# Tissue-specific Enrichment of Lymphoma Risk Loci in Regulatory Elements

**DOI:** 10.1101/017657

**Authors:** James E. Hayes, Gosia Trynka, Joseph Vijai, Kenneth Offit, Soumya Raychaudhuri, Robert J. Klein

## Abstract

Though numerous polymorphisms have been associated with risk of developing lymphoma, how these variants function to promote tumorigenesis is poorly understood. Here, we report that lymphoma risk SNPs, especially in the non-Hodgkin’s lymphoma subtype chronic lymphocytic leukemia, are significantly enriched for co-localization with epigenetic marks of active gene regulation. These enrichments were seen in a lymphoid-specific manner for numerous ENCODE datasets, including DNase-hypersensitivity as well as multiple segmentation-defined enhancer regions. Furthermore, we identify putatively functional SNPs that are both in regulatory elements in lymphocytes and are associated with gene expression changes in blood. We developed an algorithm, UES, that uses a Monte Carlo simulation approach to calculate the enrichment of previously identified risk SNPs in various functional elements. This multiscale approach integrating multiple datasets helps disentangle the underlying biology of lymphoma, and more broadly, is generally applicable to GWAS results from other diseases as well.

## Introduction

Lymphoma, including the non-Hodgkin’s lymphoma subtype chronic lymphocytic leukemia, was responsible for more than 130,000 new cases of cancer and 44,000 deaths in 2014 [1]. In an effort to understand the etiology of these diseases, numerous genome-wide association studies (GWAS) have been performed and have identified common genetic variants associated with the risk of developing lymphoma [2-14].

However, as with GWAS of other diseases, neither the identified GWAS hit nor correlated variants in linkage disequilibrium (LD) alter the amino acid sequence of a protein. We have previously observed that single nucleotide polymorphisms (SNPs) found in evolutionary conserved regions and in regions epigenetically marked for transcriptional regulation are more likely to be under negative selection in humans, suggesting biological function [15]. Others have shown that risk variants are enriched in particular epigenomic marks of transcriptional regulatory regions[16, 17] and that trait-associated SNPs, including GWAS-identified risk SNPs, are often found in expression quantitative trait loci (eQTL) that affect nearby gene expression[18]. Furthermore, recent studies have shown the etiologic nature of transcription factors themselves in some diseases[19]. Taken together, these data suggest the hypothesis that for many GWAS-identified risk lock, the functional variant may modulate disease risk through alteration of gene regulation rather than coding sequence.

To test the hypothesis that lymphoma risk SNPs, or their LD partners, tend to alter regulatory elements, we interrogated the functional genomics data from ENCODE (<u>Ency</u>clopedia <u>o</u>f <u>D</u>NA <u>E</u>lements) [20, 21]. The large amount of data available for the lymphoblastoid cell line GM12878 and other hematologically derived cells allows integrative analysis to give more accurate representation of the segments of the genome that are active regulatory elements[22, 23]. To test our hypothesis, we developed a computational pipeline, UES (<u>U</u>ncovering <u>E</u>nrichment through <u>S</u>imulation), that uses a Monte Carlo approach to test whether a set of SNPs is significantly enriched for a particular functional genomic annotation of the genome, taking linkage disequilibrium patterns into account. We demonstrate a significant enrichment of these lymphoma risk SNPs in regulatory marks specific for lymphoid tissue.

## Results

### Enrichment of CLL & Lymphoma Risk SNPs in GM12878 Regulatory Tracks

We first asked if lymphoma risk SNPs are enriched in regions annotated as putatively regulatory in GM12878 using our novel method, UES (Fig. 1). Using the NHGRI GWAS catalog [24], we identified 55 risk SNPs for lymphoma, including both the Hodgkin’s lymphoma (HD) and non-Hodgkin’s lymphoma (NHL) types. Once the list was pruned to ensure the SNPs were independent and the HLA region was excluded, the resultant list contained 36 risk SNPs (S1 Table) [2-14]. We confirmed that the minor allele distribution of our random SNPs were similar to the original input (real MAF mean = 0.278; random SNPs mean = 0.260; p=0.203). We queried the ENCODE “unified DNase” track for GM12878, which identifies regions of open chromatin regardless of the particular factors that bind. The lymphoma risk-SNPs were significantly enriched in GM12878 DNase hypersensitivity sites (p < 0.0001), with 16 distinct regions containing risk SNPs potentially explainable by a variant in a DNase hypersensitive site. The 10,000 control sets of randomly selected SNPs with similar characteristics only showed an average of 4.5 regions potentially explainable by variants overlapping a DNase hypersensitive site (Fig. 2a & S2 Table).

**Fig. 1.**
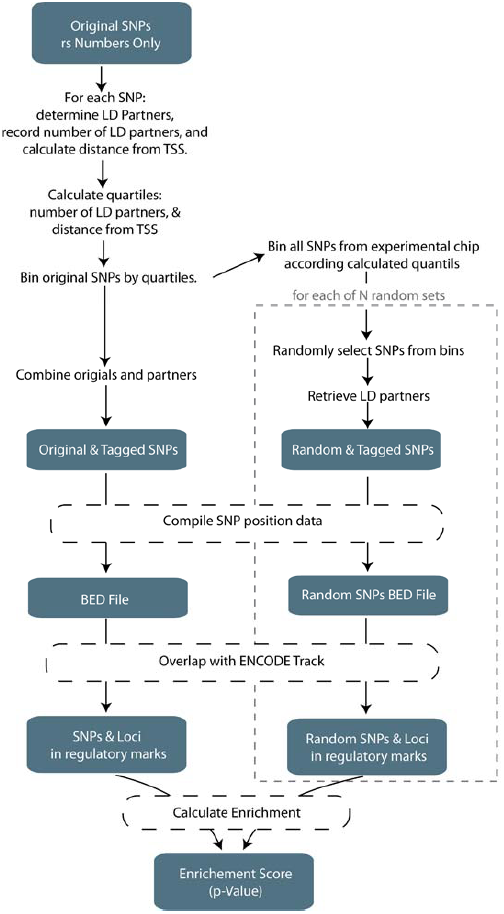
UES algorithm visualization. This represents the generalized workflow to determine the SNP enrichment in an ENCODE track. A full description and details of the algorithm can be found in the Materials and Methods.

**Fig. 2.**
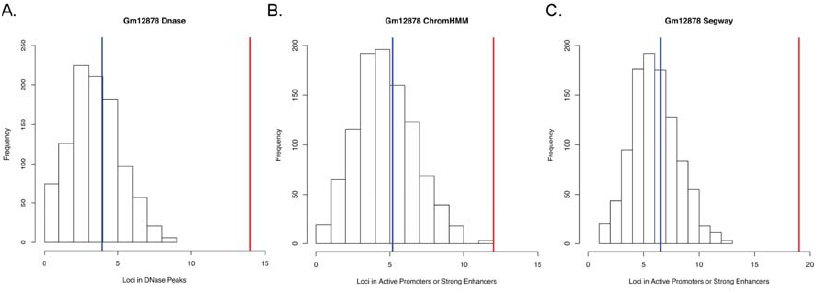
Overlap of lymphoma risk SNPs with regulatory regions in GM12878. The histograms represent the distribution of how many random loci overlap a specific annotation. The blue represents the mean of the empirical null distribution while the red line represents the real number of loci from the lymphoma and CLL GWAS that overlap the specific regulatory annotation. A, Overlap of SNPs with DNase hypersensitivity regions in GM12878. B, Overlap of SNPs with active promoters and strong enhancers as annotated by ChromHMM in GM12878. C, Overlap of SNPs with active promoters and strong enhancers as annotated by Segway in GM12878.

As some physical regions of the genome harbor more than one independent risk SNP, we were concerned this could lead to oversampling of a given region and false positives. To test this, we location pruned our input SNPs, ensuring that none of the SNPs tested were within one megabase of one another, reducing the input set from 36 initial SNPs down to 30 SNPs. We observed nearly identical results between the location pruned dataset and the original dataset (S2 Table) and concluded that these 6 extra SNPs were not falsely inflating the observed statistical result.

We next asked if enrichment could be observed in regulatory elements predicted by genome segmentation of integrated functional genomics data. Using the results from two different segmentation algorithms – ChromHMM[22] and Segway[23] – on GM12878, we asked if lymphoma risk SNPs are enriched in regions identified as active promoters or strong enhancers. We observed a significant enrichment of the lymphoma SNPs in regulatory regions as defined by both ChromHMM and Segway with p=0.0002 and p<0.0001, respectively (Figs. 2b-2c). Upon looking deeper at the ChromHMM data for GM12878, we observed the risk SNPs were enriched in each of the four classes of enhancers (2 strong enhancer classes and 2 weak enhancer classes) with p<=0.0002 when analyzed separately. When combined into a separate strong enhancer set and a weak enhancer set, we saw a significant enrichment (p<0.0001) when compared to random controls for both (S3 Table). Interestingly, the “Active Promoter” state, by itself showed no significant enrichment (p=0.3845). We observed similar results when performing the enrichment analysis for the Segway segmentation track of GM12878 (S3 Table): strong enhancers were the most enriched (p < 0.0001); weak enhancers and active promoters did not achieve significance at the Bonferoni threshold (p = 0.0031 and p=0.0419, respectively).

We hypothesized that functional SNPs may be those localized at transcription factor binding sites (TFBS). Using the same SNPs, we interrogated the set of ENCODE ChIP-Seq data for GM12878 (January 2011 data freeze). We created a master dataset consisting of the union of 75 GM12878 transcription factor ChIP-Seq data and observed a significant enrichment of the lymphoma SNPs in the peaks when compared to the random controls (p < 0.0001; S4 Table). We created additional sets containing the union of all of the transcription factor peaks with the Gm12878 DNase hypersensitivity and a union of all the LCL DNase hypersensitivity and observed similar enrichment in both (p < 0.0001, S4 Table). In order to identify specific transcription factors that bind near lymphoma risk SNPs, we ran the enrichment analysis pipeline for each factor in the ChIP-Seq dataset, 4 of which reached the significance threshold once corrected for multiple testing: NFIC, RUNX3, NF-kB, and TBLR1 (p< 0.0001; p<0.0001; p=0.0002; p=0.0005; S4 Table).

To test the validity of our findings and verify our approach, we performed the same analysis with a set of breast cancer risk SNPs identified from the NHGRI GWAS catalogue as there were a similar number of SNPs in the database at the time (n=31). Our hypothesis is that since the diseases do not share a tissue of origin, there should be no enrichment of the breast cancer SNPs in the GM12878 annotations. We did not observe statistical enrichment (random n=10,000) of the breast cancer risk loci in DNase hypersensitivity, ChromHMM enhancer, Segway enhancer data, or TF union data for GM12878 (S5 Table). We expanded our scope to see whether or not these tissue-specific observations held true when we used a different disease, prostate cancer, with roughly double the number of input SNPs (n=62). As seen with the breast cancer SNPs, there was no statistical enrichment of the prostate cancer SNPs in any of the datasets for GM12878 after correcting for multiple testing (S5 Table).

### Tissue Specificity of CLL & Lymphoma Risk SNPs

We next asked if the observed enrichments were specific to cells of the lymphoid lineage. First, we interrogated the other 124 unified DNase tracks from ENCODE with the same GWAS and random SNP sets used for the GM12878 analysis and observed enrichment of 8 additional cell lines that achieved sub-Bonferoni significance with p<0.0004. Interestingly, all of the lines that showed enrichment at that level were of the lymphoid lineage (Figs. 3a-3e, 3j-3k, S2 Table). When we relax the stringency and expand the scope to any cell lines with p<0.01, we see that 15 out of the 18 cell lines which surpassed that threshold were of the lymphoid lineage. All of the LCLs in the ENCODE database were below this threshold and had a p<=0.0065 (Figs. 3f-3i). There was only one cell line, HRE (renal epithelial cells), which was not from the lymphoid lineage that almost met the statistical threshold once corrected for multiple testing (p=0.0004). These data show that the previously reported lymphoma risk SNPs are enriched in DNase hypersensitivity regions in a tissue-specific manner (Fig. 3l).

**Fig. 3.**
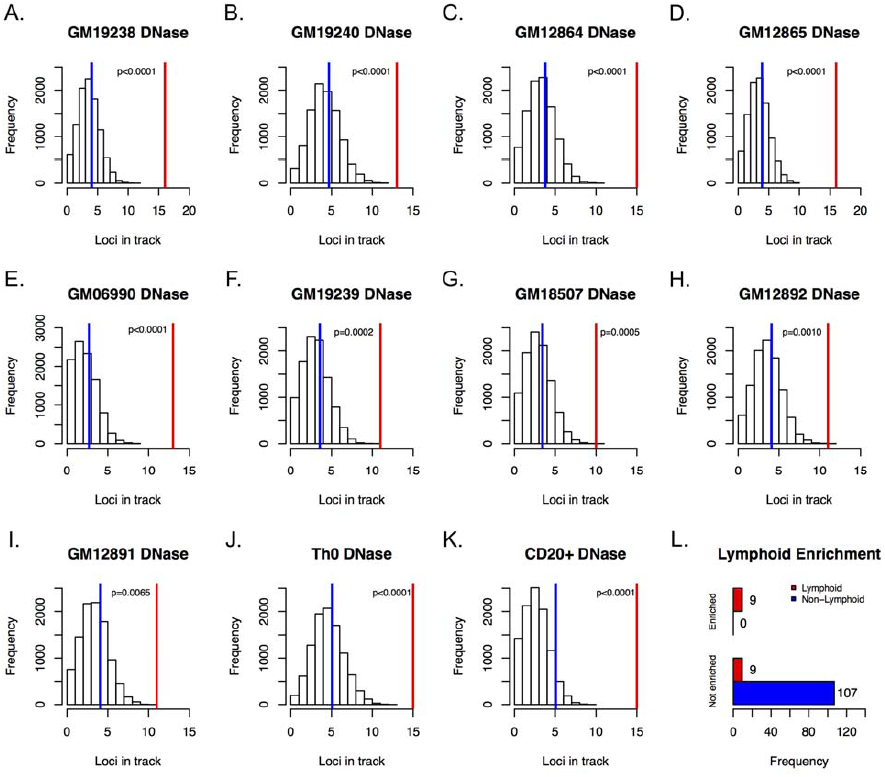
Enrichment of lymphoma and CLL risk SNPs in DNase-hypersensitive sites of lymphoblastoid cell lines. (A-I) These histograms represent the distribution of how many random loci overlap a specific annotation. The blue represents the mean of the empirical null distribution while the red line represents the real number of loci from the lymphoma and CLL GWAS that overlap the DNase hypersensitive site in the specified cell line: (A) GM19238 (B) GM19240 (C) GM12864 (D) GM12865 (E) GM06990 (F) GM19239 (G) GM18507 (H) GM12892 (I) GM12891. (J) Th0 (K) CD20+ (L) Summary of distribution of tissue of origin for cell lines in which lymphoma and CLL risk SNPs are either enriched sub-Bonferoni (p<0.0004) in DNase hypersensitive sites or not enriched.

We next performed a similar analysis on the chromatin segmentation data. We analyzed eight additional cell lines with ChromHMM segmentation data and five additional lines with Segway segmentation data for enhancer classes. For both the ChromHMM and Segway strong enhancer classifications, GM12878 was the only cell line that showed strong, significant enrichment with p=0.0002 and p<0.0001 for each dataset, respectively (S6 Table). When looking at the weak enhancer classifications, again, GM12878 was the only cell type to demonstrate any significance with p<0.0001 and p=0.0031 for the ChromHMM and Segway data, respectively (S6 Tables).

### Estimating enrichment by local annotation shifts

It has been previously noted that, under some models in which the functional variants underlying GWAS are not regulatory, enrichment of GWAS-identified SNPs in regulatory regions could occur if proper controls are not used. While our method controls for the major factors that need to be controlled for (LD patterns and distance from transcription start), we nevertheless asked if a similar enrichment could be observed with an alternative approach, GoShifter, that shifts annotations at the associated loci to test the significance of enrichment. This approach identified 5 cell lines that showed enrichment for the risk SNPs. Notably, 4 of these 5 lines that showed enrichment at p < 0.05– GM19238, Th0, Cd20, and GM06990 – are from the lymphoid lineage (S7 Table).

### Lymphoma & CLL SNPs as eQTLs

Another prediction of our hypothesis that lymphoma risk SNPs alter regulatory regions is that these SNPs will be associated with expression changes in nearby genes. To test this hypothesis, we asked how may of the published lymphoma risk SNPs are expression quantitative trait loci (eQTLs) using a recently published set of eQTLs in blood[25]. We observed that 21 of the original loci have at least 1 cis eQTL, (S8 Table). One example of the power of this overall approach is evidenced by rs7097. This SNP was initially defined as a lymphoma risk SNP[8] but did not intersect with any LCL DNase hypersensitivity sites, nor with promoter or enhancer regions of chromatin segmentation data. However, one of the SNPs it tags, rs694609, was found in a DNase hypersensitivity site of GM12878, which was categorized as falling in an “Active Promoter” by ChromHMM or a “Transcription Start Site” by Segway. Even as both the original SNP, rs7097, and the tag, rs694609, showed evidence of being *cis*-eQTLs for the same genes (POLR1D, LNX2, and GTF3A), our pipeline would suggest that the tagged SNP is the more likely candidate to be the functionally relevant SNP as it is found in open chromatin and in functional marks (S8 Table).

## Discussion

We have shown significant enrichment of previously reported lymphoma and CLL risk SNPs in regulatory elements in lymphoblastoid cell lines, pointing to the tissue specific manner through which these genetic loci may confer increased risk for lymphomagenesis. Looking specifically at the analyses in GM12878, we saw this enrichment in DNase hypersensitivity loci as well as numerous enhancer sites; we did not observe enrichment when we looked at risk loci for other cancer types. We have also identified candidate functional SNPs that co-localize with these genomic marks and have also been shown to be eQTLs in published blood datasets.

We were able to perform these analyses because there are significant amounts of functional genomic data available for the cell line GM12878. Taking a deeper at those results, we note a similar number of loci for which a candidate functional SNP can be found in DNase regions, ChrommHMM-Strong Enhancers, and Segway-Strong Enhancers (n=16, 12, and 17, respectively). While none of these 3 datasets were complete subsets of each other, there is significant overlap (S8 Table). However, as DNase hypersensitivity data can be obtained from a single assay as opposed to a combination of multiple assays for the segmentation data, in the case where data on relevant cell types do not yet exist, DNase data may be sufficient to identify putatively functional SNPs before investing the time and resources to generate all the assays needed for segmentation analysis.

We believe that these enrichment studies can provide valuable insight into the potential etiology of the disease of interest. For example, looking at the enrichment of lymphoma SNPs in ChIP-Seq data, we see an enrichment of risk SNPs in RUNX3 binding sites(p<0.0001). *RUNX3* is a gene which is highly expressed in LCLs [26] and has been shown, paradoxically, to act both in promoting and suppressing tumor growth [27]. We also observed enrichment of risk SNPs in binding sites for Nf-kB and TNF (p<0.0001); variation within these two pathways have also been shown to associate with non-Hodgkin’s lymphoma risk [28]. Lymphoma risk SNPs are also enriched at binding sites of *TBLR1* (p=0.0005); disruptions at the *TBLR1* locus in diffuse large B-cell lymphoma have been seen through a deletion of the locus [29] and the identification of a novel fusion between it and *TP63*, a paralogue of *TP53* [30].

We acknowledge that there are pipelines similar to our UES algorithm that are already deployed and perform in similar ways. However, the UES pipeline makes improvements on those methods through the controlling for the input SNPs based on the initial SNP chip the study was done on, the distance a SNP is from a TSS, and the number of LD partners to compute an empirical p-value. While RegulomeDB[31] and HaploReg[32] both work quickly, thoroughly, and robustly annotate the input SNPs with genomic regulatory data they do not provide a formal statistical test of enrichment. Though Maurano et al.[16] were able to show a distinct enrichment of GWAS SNPs in DNase hypersensitivity sites, by accounting for the particular genotyping platform used in a GWAS we reduce the risk of spurious enrichment signals due to a nonrandom distribution of SNPs from the GWAS platform with respect to genomic features. Trynka et al.[17] also used a permutation approach to test for significance of enrichment, but their approach focuses on the question of tissue specificity rather than a general test of enrichment.

More recently, Trynka et al. [33] discuss the pitfalls of a SNP-matching approach which lead to an over-inflation of the significance scores. However, their study also demonstrates that a large amount of the over-inflation of results can be corrected when taking into account the LD structure of the input SNPs, which UES does. Both approaches agree with each other and show significant enrichment of the lymphoma SNPs in the DNase hypersensitivity sites for numerous LCLs. While acknowledging the strength of GoShifter, UES provides an improvement over the other SNP-matching algorithms in the field today and can provide useful insights that were not captured by GoShifter.

Overall, the data presented support the hypothesis that regulatory variants that influence transcription in cells of the lymphoid lineage contribute to inherited risk of lymphoma and chronic lymphocytic leukemia. These results validate our computational approach that, moving forward, could provide novel insight into disease etiology when applied to other diseases.

## Methods

### Pipeline Construction and Workflow

We developed a computational pipeline entitled “Uncovering Enrichment through Simulation” (UES) to test if GWAS-identified SNPs are enriched in particular functional annotations through use of Monte Carlo simulations. The pipeline (Fig. 1) is written predominantly in Perl and accepts 3 parameters: a text file containing the input set of SNPs, the genotyping platform from which to choose the random sets, and the number of random sets to be constructed. SNPs that had been identified at the HLA region – defined as chr6:29570005-33377658 (build 37) – were removed due to the high amount of variability and linkage disequilibrium at that region. Each of the initial SNPs is then categorized by its distance from the nearest transcription start site (TSS) and its number of LD partners. Quartiles for both the TSS distance and LD partner count are calculated separately, and the initial SNPs are binned accordingly. The number of each of the initial SNPs contained in each bin (characterized by distance from TSS and LD partner count) is recorded and used for subsequent random SNP set selection. Upon completion of this step, all of the SNPs from the appropriate genotyping platform are loaded (excluding the HLA region) and binned according to the initial SNP criteria. Since it has been shown that disease-associated SNPs have a higher MAF than expected by chance[34], we filter the platform SNPs and keep only those with a MAF >= 5% keeping in concert with the common filter steps when performing GWAS. Random SNP sets are chosen, matching the original bin frequencies, and LD partners are retrieved (r^2^>0.8). All the data have been pre-calculated and are retrieved using Tabix[35]. The script executes an instance of BedTool’s intersectBed[36] in order to determine which SNPs fall directly in a given track. Those resultant SNPs are then collapsed into loci that co-localize with marks based on LD structure. Finally, the empirical p-value for a specific track is calculated by the following formula:

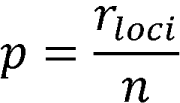

where *r*_*loci*_ = the number of instances when the frequency of co-localization of the random SNP sets with the feature >= the number of loci that co-localize with the feature for the initial input set of SNPs, and *n* = the number of random-SNP sets chosen. The current pipeline and subsequent versions are available for download from the Klein lab’s website http://research.mssm.edu/kleinlab/software/ues. Algorithm development can be found at

### CLL & Lymphoma Risk SNPs

We manually queried the NHGRI GWAS Catalog and selected a master list of CLL/lymphoma SNPs that had been reported as having a significant association. To ensure independence, for any SNPs that were correlated (r^2^ > 0.8), the SNP with the lower reported p-value was kept. Initially, 56 CLL & lymphoma SNPs were entered into the pipeline, and once the HLA region was excluded, there were 36 SNPs used for the remainder of the analysis (S1 Table). Next the LD partners were found, resulting in 591 SNPs used for analysis of the original lymphoma and CLL data. The enrichment pipeline produced 10,000 sets consisting of 36 matched random SNPs. Once LD partners were included, the sets used for analysis range in size from 331 to 4028 SNPs.

### Location Pruning of CLL & Lymphoma Risk SNPs

In order to ensure that the observed signal was not due to oversampling of a region, we pruned SNPs from the input set so that SNPs were separated by at least a megabase. For those SNPs in close proximity, we retained the SNP that had the lowest reported p-value, resulting in a set of 30 input SNPs.

### Pipeline Run Parameters

Since our analysis was run on a collection of SNPs from multiple studies, we provided the parameter that chose the matched-random SNPs from a union set of both Illumina and Affymetrix genotyping chips. The pipeline outputted 10,000 sets of feature-matched random SNPs.

### Regulatory Track Data

The ENCODE datasets were obtained directly from the ENCODE Consortium’s website. The DNase hypersensitivity analysis was performed using the ENCODE Consortium’s “unified DNase hypersensitivity” tracks (http://hgdownload-test.cse.ucsc.edu/goldenPath/hg19/encodeDCC/wgEncodeAwgDnaseUniform/). The ChromHMM track was also downloaded from ENCODE (http://hgdownload-test.cse.ucsc.edu/goldenPath/hg19/encodeDCC/wgEncodeAwgSegmentation/), after which a Perl script was used to extract the active promoter, strong enhancer, and weak enhancer regions, or combine the active promoter and strong enhancer regions into a combination track. The Segway segmentation was downloaded directly from the Noble lab’s website and was modified in the same way as described for the ChromHMM data (http://noble.gs.washington.edu/proj/segway/).

### Peak-Shifting (GoShifter)

We also used the new software GoShifter to test for enrichment. Using European samples from the 1000 Genomes dataset we first iterated over each locus to identify all the variants in tight LD (r^2^>0.8). Locus boundaries were defined by the most downstream and upstream LD SNP and extended by two times the median size of a tested annotation. We then circularized the locus and allowed annotations to randomly shift in 10,000 iterations. For each of the shifting iterations we quantified the number of loci at which a variant overlapped with an annotation. The reported p-value corresponds to the number of iterations where enrichment exceeded the observed value[33].

## Acknowledgements

This work was supported in part through the computational resources and staff expertise provided by the Department of Scientific Computing at the Icahn School of Medicine at Mount Sinai. We are grateful to Mridu MIddha in helping beta test the software.

## Supporting Information

**S1 Table. Lymphoma SNPs identified from the NHGRI GWAS catalog.** The number of SNPs was reduced to 36 from 55 once those SNPs at the HLA region were identified and removed.

**S2 Table. Enrichment of lymphoma SNPs in ENCODE Unified DNase tracks.** Enrichment analysis using the UES pipeline were performed for each of the 125 DNase hypersensitivity tracks in the ENCODE database. Cell line information (Tissue, Blood cell classification, and karyotype) were obtained directed from ENCODE’s description of the cells used. The “OrigLoci” column gives the number of loci (once the SNPs are collapsed into loci based on LD partners) for the input lymphoma & CLL SNPs that overlapped with the specific mark. The “Rand>=Orig” column is the number of times a random SNP file had greater than or equal to the number of loci co-localizing with the particular mark. The “Random_Avg” column is the average of the 10,000 random generated SNP sets and the loci that overlap with the mark. The “pValue” is calculated by taking the number of random SNP sets that were greater than or equal to the input SNPs divided by n, in this case, 10,000. The “location pruned pValue” is the reported p-value for the rerun of the analysis using the input data where SNPs were removed within one megabase of one another. (See Methods.)

**S3. Table. Enrichment of lymphoma risk SNPs in GM12878 segmentation data.** We ran the UES pipeline for each of the segmentation tracks, both ChromHMM and Segway, for GM12878. These data are calculated and represented in the same way as Supplementary Table S2.

**S4. Table. Enrichment of lymphoma risk SNPs in GM12878 Chip-Seq data.** The UES pipeline was run to calculate the enrichment of the lymphoma risk SNPs with with transcription factor binding data in GM12878. The “Gm12878 ChipSeq” Union track was created by merging all of the ChipSeq tracks for GM12878. That track was subsequently merged with the DNase hypersensitivity track of GM12878 to create the “Gm12878 DNase & Gm12878 ChipSeq Union”. The “All LCL Union” track was constructed in a similar manner by merging all of the DNase hypersensitivity tracks for the 10 LCLs in the ENCODE database. The “All LCL Union & Gm12878 ChipSeq Union” track was created by merging both of those union tracks. These data are calculated and represented in the same way as Supplemental Table S2. This table also reports the p-values for the analysis of the individual transcription factor data for GM12878. The column descriptions are the same as in Supplementary Table S2.

**S5. Table. Enrichment analysis of breast cancer and prostate cancer risk loci in GM12878 segmentation data.** These data are calculated and represented in the same way as Supplementary Table S2.

**S6 Table. Enrichment of lymphoma risk SNPs in “Active Promoter & Strong Enhancer,” “Strong Enhancer,” and “Weak Enhancer” segmentation datasets for both ChromHMM and Segway.** “Strong Enhancer” tracks (combination tracks made of strong enhancer states 4 & 5) for all the ChromHMM cell types. These data are calculated and represented in the same way as Supplementary Table S2.

**S7 Table. GoShifter enrichment of lymphoma risk SNPs in DNase hypersensitivity data.**

**S8 Table. Summary of lymphoma risk SNPs.** Summary of all original lymphoma & CLL SNPs and their LD partners, along with information on whether or not any of these SNPs were found to be in DNase hypersensitivity sites, active promoters (for both ChromHMM and Segway), or strong enhancers (ChromHMM and Segway). All of the tracks are for GM12878. The final column describes the eQTL status of the as seen in Westra et al.

